# Revisiting alpha-theta cross-frequency dynamics during working memory

**DOI:** 10.1101/2025.08.18.670850

**Authors:** Julio Rodriguez-Larios, Roberts Mark J., Saskia Haegens

## Abstract

Prior Electroencephalography (EEG) research has shown that during working memory delay, alpha (8–14 Hz) and theta (4–8 Hz) oscillations tend to form a 2:1 frequency ratio. According to the Binary Hierarchy Brain Body Oscillation Theory (BHBBOT), a recent model grounded in mathematical analysis, such harmonic (2:1) alpha:theta frequency configurations reflect enhanced connectivity between brain regions generating these rhythms. However, this prediction has not yet been empirically tested. In this study, we leveraged Information Theory and the Theory of Weakly Coupled Oscillators (TWCO) to examine whether the previously observed frequency modulations in alpha and theta rhythms during working memory are accompanied by changes in inter-areal connectivity. Contrary to the BHBBOT predictions, both Information Theory metrics and TWCO parameters showed that connectivity between frontal theta and parietal alpha rhythms was significantly reduced during the working-memory delay period (while the proportion of 2:1 ratios increased). In addition, phase locking value, a standard measure of synchrony, was also significantly reduced during working memory delay and was negatively associated with behavioural performance. In conclusion, our results show that the increased occurrence of 2:1 alpha:theta cross-frequency ratios during working memory reflects functional segregation (rather than integration) between frontal and parietal regions.

## Introduction

Cognitive functions require the coordinated activity of many brain areas (Christophel et al., 2017), and neural oscillations are thought to support this (X.-J. Wang, 2010). Brain oscillations are rhythmic patterns of neural activity that create sequences of excitation and inhibition (Buzsáki & Draguhn, 2004). It has been proposed that proposed that neural oscillations gate inter-areal communication by the alignment of excitatory phases (or ‘duty cycles’) that make both spike output and sensitivity to synaptic input coincide in time (Fries, 2015; Varela et al., 2001). In support of this idea, phase locking between areas oscillating at a similar frequency has been shown to be behaviourally relevant in both human and animal neurophysiology (Spyropoulos et al., 2018; Vissani et al., 2025).

Different brain areas tend to generate oscillations at different frequencies (Capilla et al., 2021), depending on the neurophysiological characteristics of their neural populations (Buzsáki & Draguhn, 2004; Lea-Carnall et al., 2016; von Stein & Sarnthein, 2000). The interaction between areas that oscillate at different frequencies is thought to be captured by various cross-frequency coupling metrics (e.g., phase-phase, phase-amplitude, amplitude-amplitude) (Canolty & Knight, 2010; Palva & Palva, 2017). According to the Binary Hierarchy Brain Body Oscillation Theory (BHBBOT), areas that oscillate at different frequencies can transiently couple by forming harmonic cross-frequency ratios (e.g., 2:1, 3:1, 4:1, etc.) and decouple by forming non-harmonic irrational ratios (e.g., 1.6:1) (Klimesch, 2013, 2018; Rassi et al., 2019). This prediction is based on mathematical analysis showing that while harmonic ratios would lead to stable phase relationships and therefore enhance “coupling”, those given by irrational numbers (such as the golden mean) would lead to variable phase relationships thereby facilitating a “decoupled state” (Pletzer et al., 2010).

Cross-frequency ratios between alpha and theta rhythms have been investigated in relation to different mental states and cognitive functions (Rodriguez-Larios, Faber, et al., 2020; Rodriguez-Larios, Wong, et al., 2020; Rodriguez-Larios & Alaerts, 2019; Rodriguez-Larios & Alaerts, 2020). In the context of working memory, alpha and theta rhythms showed an increased occurrence of cross-frequency ratios around 2:1 and a decreased occurrence of cross-frequency ratios below 1.7:1 during memory retention(Rodriguez-Larios & Alaerts, 2019). This was in line with previous literature showing increases in alpha frequency (Haegens et al., 2014; Mierau et al., 2017) and decreases in theta frequency (Senoussi et al., 2022) during working-memory delays. Although according to the BHBBOT (Klimesch, 2018), the harmonic (2:1) frequency configuration of alpha and theta rhythms during working memory would lead to greater coupling between the brain areas generating these rhythms, this prediction has not been empirically tested.

Information theory provides a model-free framework for quantifying (non-linear) statistical dependencies between different brain regions (Fagerholm et al., 2023; Vergara et al., 2017). Among the most used metrics in neuroscience are mutual information (MI) and transfer entropy (TE) (Timme & Lapish, 2018). MI quantifies the shared information between two signals and is typically used to assess functional connectivity, while TE captures the directed, time-lagged influence of one signal on another, making it suitable for assessing effective connectivity (Lindner et al., 2011; Vicente et al., 2011). Both of these metrics are based on the concept of entropy, which measures the level of uncertainty or unpredictability associated with a random variable (Fagerholm et al., 2023; Lu & Rodriguez-Larios, 2022). MI uses entropy to quantify the reduction in uncertainty about one signal given knowledge of another, effectively measuring how much information is shared between two signals. TE extends this by quantifying how much the past state of one signal reduces the uncertainty of the future state of another signal, conditioned on the target’s own past—thereby capturing directional, time-lagged interactions.

The Theory of Weakly Coupled Oscillators (TWCO) offers a mechanistic framework for understanding brain connectivity based on transient phase synchrony between oscillators with different natural frequencies (Breakspear et al., 2010; Ermentrout & Kleinfeld, 2001; Kuramoto, 1991). In this view, synchronisation is a non-stationary and non-linear process that, based on frequency modulations, allow groups of oscillators to coordinate around a preferred phase-relation (Lowet et al., 2022). This theory predicts that synchrony between oscillators depends on the interaction between two parameters: coupling strength and frequency detuning (Hoppensteadt & Izhikevich, 1998; Lowet et al., 2022). While coupling strength reflects phase adjustments, detuning is equivalent to mean frequency difference (or numerical ratio) between oscillators. Previous work has shown that gamma band oscillations originating from neural populations in monkey visual cortex behave like weakly coupled oscillators (Lowet et al., 2017). Based on these results, it was proposed that TWCO would apply to other brain areas and brain rhythms (Lowet et al., 2017, 2022).

In the present study, we aim to determine whether the previously observed shift toward 2:1 alpha:theta cross-frequency ratios during working-memory retention in EEG is associated with changes in connectivity between the regions generating these rhythms. To this end, we combine model-free metrics from information theory (MI and TE) with model-based parameters derived from TWCO. We use EEG data collected while participants performed a visual working-memory task (N = 26) and focus on frontal theta and parietal alpha components isolated via source-resolved spatio-spectral decomposition. This approach allowed us to test whether a putative increase in alpha:theta harmonicity (i.e., increased occurrence of 2:1 cross-frequency ratios) during memory retention corresponds to changes in inter-areal communication.

## Methods

This study used a publicly available EEG dataset that was originally published by Rodriguez-Larios and Haegens (2023). The dataset can be found at https://osf.io/fkhnb/ and MATLAB code of the entire analysis pipeline can be found here: https://osf.io/d9rc7/?view_only=26941f4be9744c5691863bef92b7ea9c

### Participants

The dataset comprises data from 31 healthy adult volunteers (13 male), with a mean age of 32.5 years (SD = 8.5). All participants had normal or corrected-to-normal vision and no self-reported history of neurological or psychiatric conditions. The study protocol and consent procedures were approved by the Institutional Review Board at the New York State Psychiatric Institute (Protocol Number 8001). Participants were compensated $25 per hour for their time. Due to technical issues during EEG acquisition or the presence of unresolvable EEG artefacts, 5 participants were excluded from analysis, resulting in a final sample size of 26 individuals.

### Design and Task

Participants engaged in a visual working-memory task (**Figure 1**) while EEG was recorded. Each trial began with a 1 second fixation cross. This was followed by the presentation of one or three oriented bars arranged in a circle, with each bar presented sequentially for 1 second each. After a 3-second retention delay, a cue appeared for 1 second, instructing participants to either retain (“stay”) or mentally reverse (“switch”) the presented orientation. Following this instruction, a response-mapping screen appeared, indicating which of the eight numbered buttons corresponded to each possible angle. This mapping changed on every trial. After participants responded, feedback was provided in the form of a green (correct) or red circle (incorrect). The full session lasted approximately one hour. For the current study, we only analysed data from Fixation and Delay periods when the load was 3, giving., a total of 96 trials per participant.

**Figure 1.**
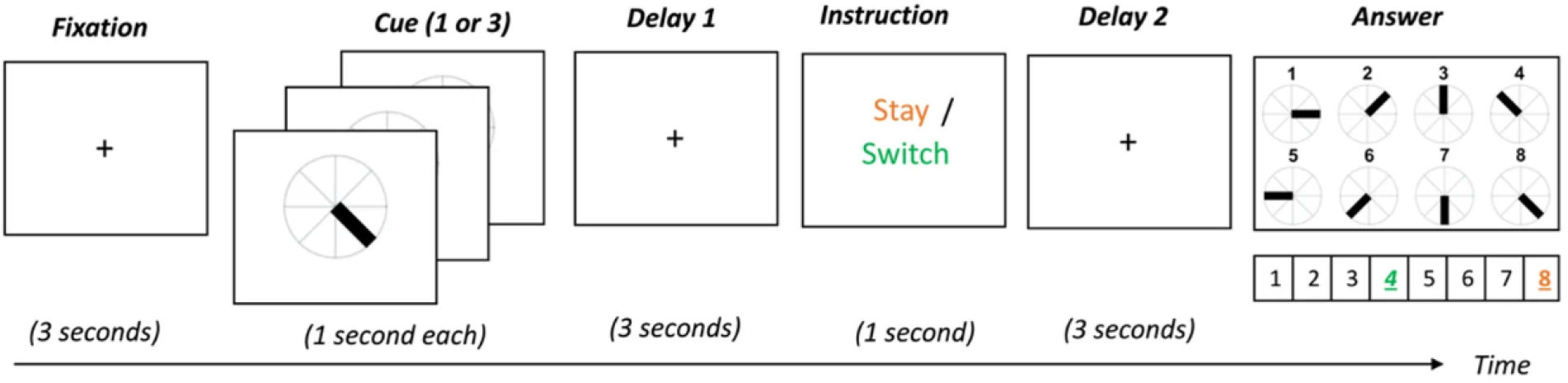
Working-memory task. In each trial, participants were shown either one or three visual stimuli and instructed to memorise their orientations (corresponding to Load 1 or Load 3, respectively). Following a cue, they were asked to indicate either the original angle (“Stay” condition) or its opposite angle (“Switch” condition) using the keyboard. In the example shown, colours represent the correct response: orange for “Stay” (angle 4) and green for “Switch” (angle 4).

### EEG acquisition and preprocessing

EEG signals were recorded from 96 scalp electrodes using the BrainVision actiCAP system (Brain Products GmbH, Munich, Germany) at a sampling rate of 500 Hz. Electrodes were placed according to the 10–20 system, with Cz used as the reference during acquisition. Signal amplification and digitisation were performed using a Brain Products actiCHamp DC amplifier connected to BrainVision Recorder software (version 2.1). Eye movements were recorded using bipolar electrodes placed vertically (above and below the left eye) and horizontally (at the outer canthi of both eyes). Additionally, electrocardiographic activity (ECG) was recorded using bipolar chest electrodes.

Data preprocessing was carried out in MATLAB R2024b using custom scripts based on EEGLAB functions (Delorme & Makeig, 2004). The data were band-pass filtered between 1 and 30 Hz and then re-referenced to the common average. Independent Component Analysis (ICA) was applied, and artefactual components were automatically identified and removed using the ICLabel algorithm (Pion-Tonachini et al., 2019) targeting components associated with muscle, ocular, cardiac, or channel noise. Any component showing an absolute correlation above 0.8 with the VEOG, HEOG, or ECG channels was also excluded. Furthermore, the Artifact Subspace Reconstruction (ASR) method was used to detect and correct transient artefacts, with a cutoff threshold set at 20 standard deviations (Chang et al., 2020).

### Separation of frontal theta and parietal alpha rhythms

Frontal theta and parietal alpha rhythms were first separated using the Spatio-Spectral Decomposition (SSD) algorithm (Nikulin et al., 2011) (see MATLAB implementation here: https://github.com/svendaehne/matlab_SPoC/blob/master/SSD/ssd.m). In short, SSD maximises the signal power at a target frequency while minimising power at neighbouring frequencies, thereby enhancing the signal-to-noise ratio (SNR) of oscillatory components. The algorithm uses a generalised eigenvalue decomposition of covariance matrices derived from band-pass and flanking frequency filters. The SSD output of is a set of spatial filters that optimally project the multichannel EEG data to extract components with maximal power at a specific frequency band and minimal power in neighbouring frequency bins.

In order to differentiate between frontal theta and parietal alpha components, the sources of SSDs were estimated using the DIPFIT plugin implemented in EEGLAB. For the identification of the midfrontal theta component, we selected the dipole with highest SNR in medial prefrontal and anterior cingulate regions (including the rostral and caudal anterior cingulate, superior frontal, medial orbitofrontal, frontal pole, and paracentral areas from the Desikan–Killiany atlas). To identify the parietal alpha component, we selected the parietal dipole (including the superior and inferior parietal lobules, precuneus, and supramarginal gyrus) with the highest SNR. **Figure 2** shows the average topography, dipole density and spectrum of frontal theta and parietal alpha components.

**Figure 2.**
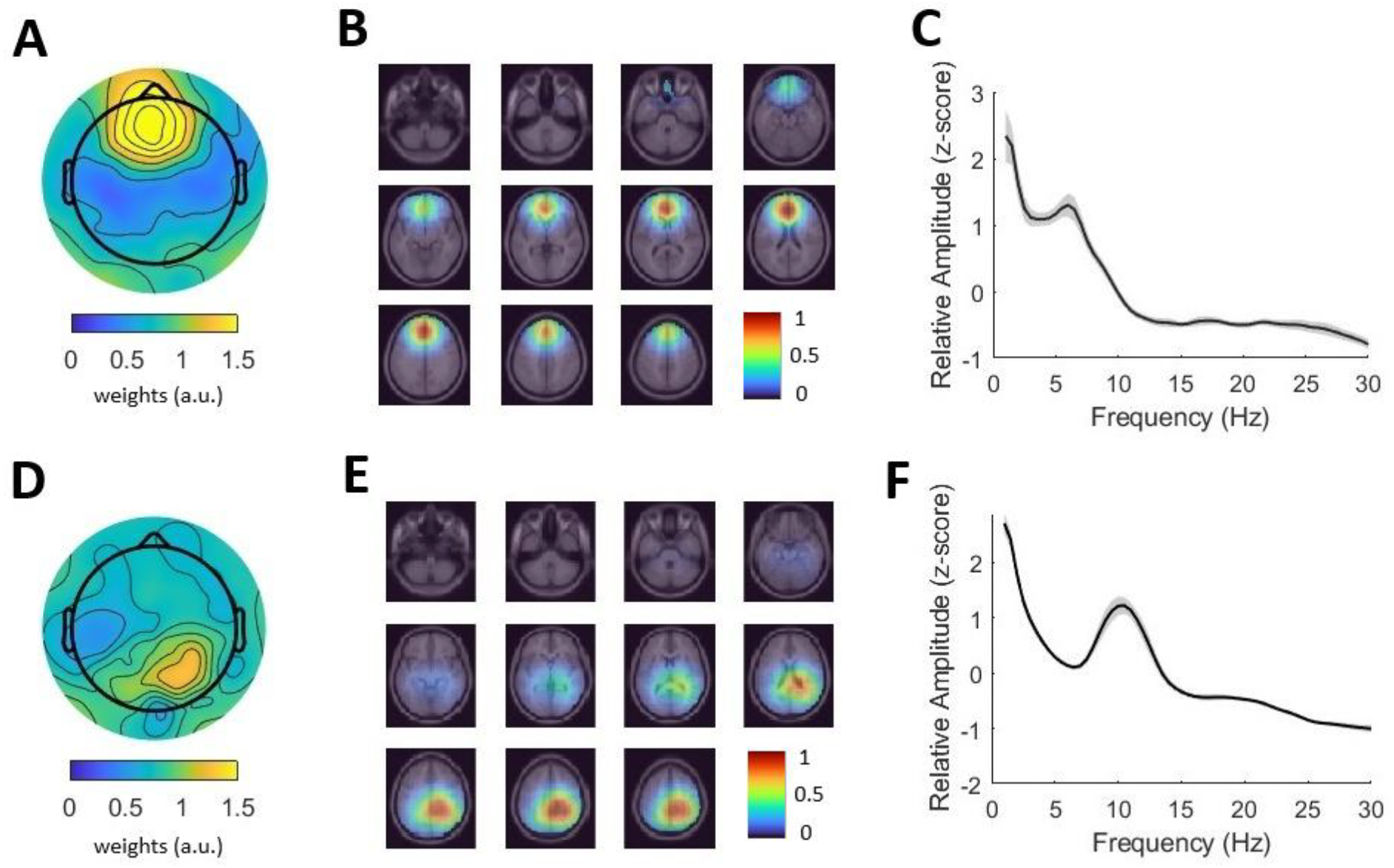
Frontal theta and parietal alpha components. **A)** Mean topography of selected frontal theta components. The absolute values of the spatial patterns given by the SSD algorithm were averaged across subjects. **B)** Dipole density of selected frontal theta components. Heatmap reflects the normalised density of dipoles in each voxel. **C)** Average spectrum of selected frontal theta components. Amplitude was normalised in the frequency dimension (z-score) per subject and component. Shaded area depicts standard error. **D)** Mean topography, **E)** dipole density and **F)** average spectrum of selected parietal alpha components. Conventions as in A to C.

### Estimation of cross-frequency ratios

Modulations in alpha:theta cross-frequency ratios were estimated as in Rodriguez-Larios & Alaerts (2019). First, data was transformed with short term Fourier transform (1-second window, 90% overlap, 0.1-Hz resolution between 1 and 30 Hz) using the MATLAB function *spectrogram*. Then, transient peaks for the theta and alpha components were detected using the MATLAB function *findpeaks*. Only peaks with amplitude above an estimate of the 1/f trend were used. This estimate of the 1/f trend was computed by linear fitting in log-log space of the spectrum per component across conditions (Rodriguez-Larios & Haegens, 2023). The proportion of different cross-frequency ratios (from 1 to 3.5) were estimated by dividing transient alpha and theta peak frequencies and rounding to the first decimal.

### Information theory metrics

Mutual information (MI) and transfer entropy (TE) between frontal theta and parietal alpha components were estimated following the procedures described by Timme and Lapish (2018). To compute MI, time series of the theta and alpha components were first discretised using a uniform count binning procedure to maximise sensitivity to interactions between the signals. MI was then computed as the reduction in uncertainty of one component given the state of the other, using the standard definition based on joint and marginal probability distributions. TE was estimated in both directions (frontal theta → parietal alpha; parietal alpha → frontal theta) using the conditional mutual information formulation of transfer entropy. A range of neurophysiologically plausible lags was explored by computing TE for delays corresponding to 1–20 samples (4–80 ms at a sampling rate of 250 Hz), as recommended to account for unknown interaction delays (Timme & Lapish, 2018). TE values were then averaged across these lags to obtain a robust measure of directed information flow. All information-theoretic analyses were implemented using the MATLAB code provided in the Neuroscience Information Theory Toolbox (https://github.com/nmtimme/Neuroscience-Information-Theory-Toolbox).

### Estimation of phase locking and TWCO parameters

To assess connectivity between frontal theta and parietal alpha components, we computed phase locking value (PLV) and the parameters coupling strength and detuning, derived from TWCO. For each component, data were first filtered using zero-phase FIR filters (4–8 Hz for frontal theta and 8– 14 Hz for parietal alpha). Then instantaneous phase and frequency were computed through the Hilbert transform, with instantaneous frequency estimated using a Savitzky-Golay derivative filter to smooth phase trajectory and avoid outliers (Lowet et al., 2017; Schafer, 2011). PLV was quantified as the mean vector length of phase differences (averaged over a sliding-window of 300-ms with 50-ms steps) within each trial. Coupling strength and detuning were estimated from the TWCO phase response curves (PRCs), constructed by binning phase differences (24 π/12 bins) and calculating median frequency differences per bin, with all data pooled across trials (Lowet et al., 2017). PRCs were smoothed with a circular moving average, and a smoothing spline was fitted to the data. Detuning was defined as the mean frequency difference across the PRC, while coupling strength was estimated as the peak-to-trough range of the fitted PRC, reflecting the amplitude of the phase– frequency interaction (see **Figure 5A** for depiction). All analyses were performed in MATLAB using custom scripts.

### Statistical analysis

Condition-related differences for each parameter were assessed using paired-samples *t*-tests (as implemented in MATLAB), with corresponding effect sizes calculated using Cohen’s *d* (Nakagawa & Cuthill, 2009). To examine the relationship between EEG-derived parameters and behavioural performance (accuracy and reaction time), a multiple linear regression analysis was conducted. Specifically, regression analyses were implemented in MATLAB using the *stepwiselm* function. This approach performs a stepwise linear regression by iteratively adding or removing predictors based on their contribution to model fit, evaluated using the Bayesian Information Criterion (BIC). Significance level for all tests was established at *p* < 0.05. Multiple comparison correction was performed through False Discovery Rate (Benjamini & Hochberg, 1995).

## Results

### Alpha:theta cross-frequency dynamics

Parietal alpha components showed a significant increase in frequency (*t*(25) = 3.08, *p* = .005, *d* = 0.60) and a significant decrease in amplitude (*t*(25) = -2.49, *p* = .020, *d* = 0.49) during the working-memory delay relative to fixation. In contrast, frontal theta components showed a significant decrease in frequency (*t*(25) = -2.38, *p* = .026, *d* = 0.47) and a significant increase in amplitude (*t*(25) = 3.55, *p* = .002, *d* = 0.70) during the working-memory delay compared to fixation (**Figure 3AB**).In terms of cross-frequency ratios, the working-memory delay showed a significant increase in ratios between 1.8 and 2.1 and at 2.4, and a decrease in ratios between 1.0 and 1.3 (*p* < .05, FDR-corrected; **Figure 3C**) relative to fixation.

**Figure 3.**
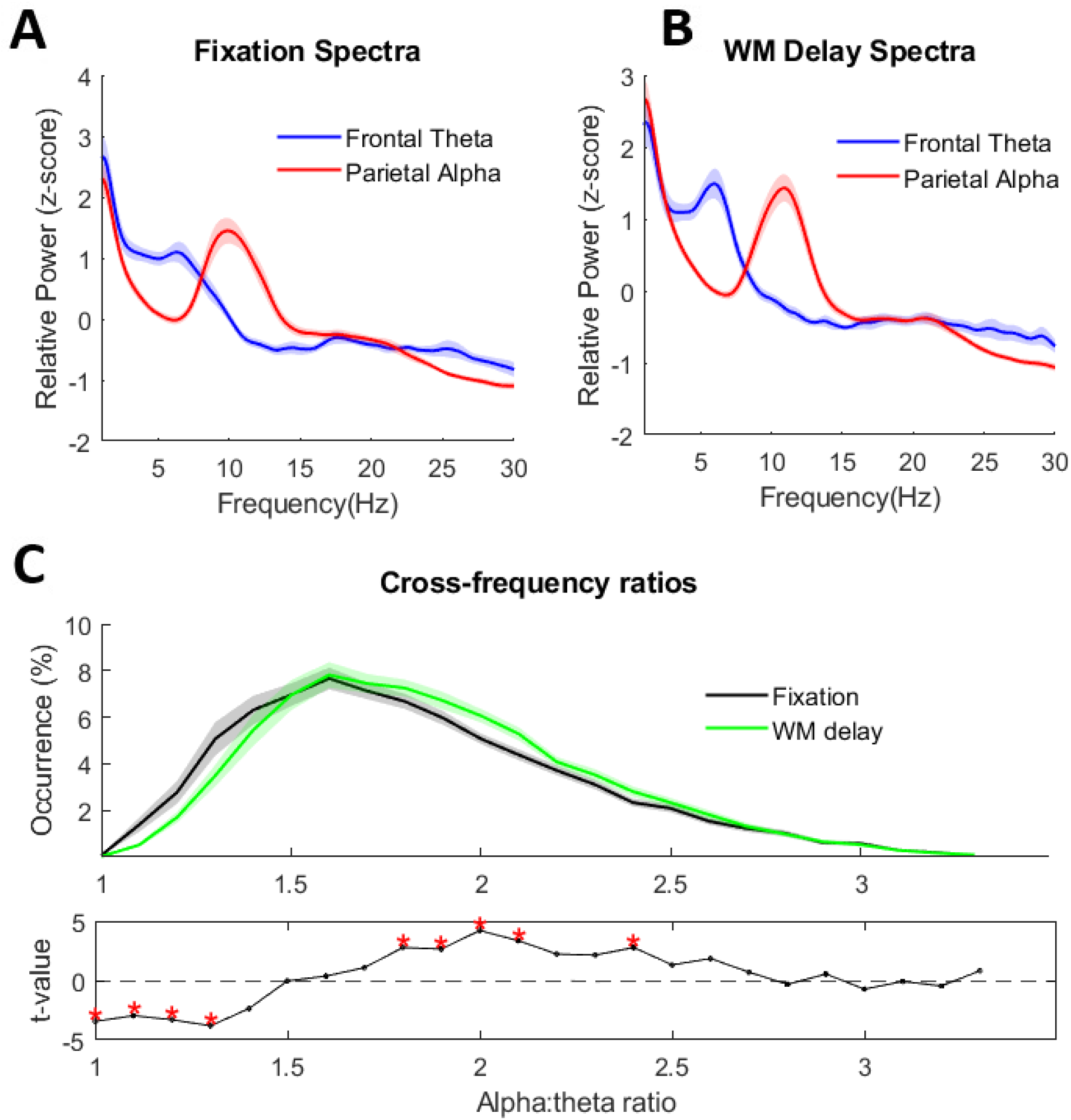
Modulations in alpha:theta cross-frequency dynamics during working memory. **A)** Power spectra of frontal theta (blue) and parietal alpha (red) components during **A)** fixation and **B)** working-memory (WM) delay. **C)** Relative occurrence of different alpha:theta cross-frequency ratios in each condition (top) and t-values resulting from the comparison (bottom). Significant ratios (p<0.05 after FDR correction) are marked with red asterisks.

### Information theory metrics

Mutual information was significantly reduced during the working memory delay compared to fixation (*t*(25) = –2.29, *p* = .031, *d* = 0.45; **Figure 4A**). Similarly, transfer entropy in the anterior-to-posterior direction was significantly reduced (*t*(25) = –3.34, *p* = .003, *d* = 0.65; **Figure 4B**), while no significant differences were found in the posterior-to-anterior direction (*t*(25) = –0.90, *p* = .377, *d* = 0.18).

**Figure 4.**
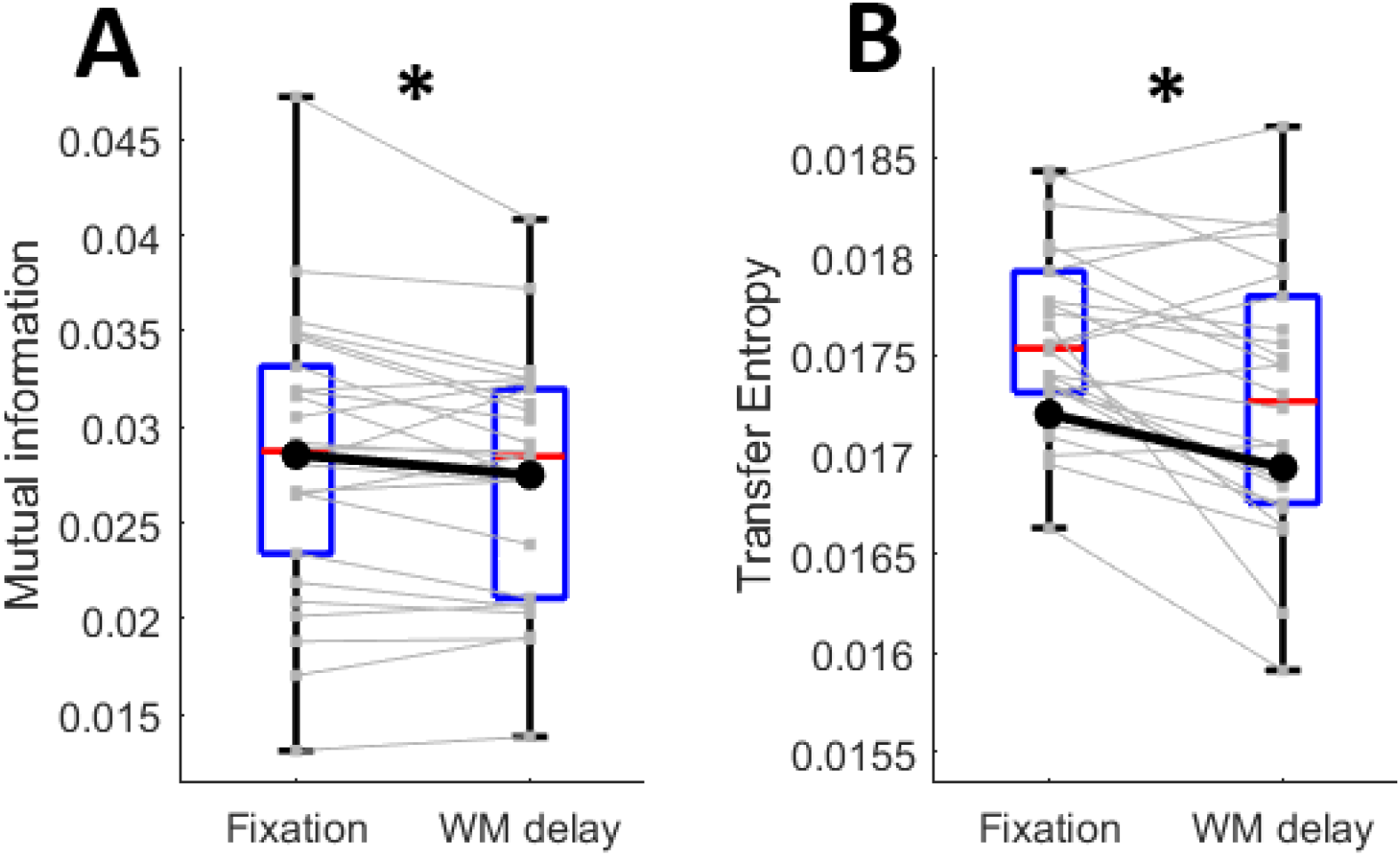
Condition-related modulations in information theory metrics. **A)** Mutual information and **B)** transfer entropy in the anterior–posterior direction were significantly reduced during working-memory delay (relative to fixation). Individual lines reflect subjects, the blue boxes indicate the 25th and 75th percentiles and red centrelines show the median in each condition. Asterisks indicate statistical significance at p<0.05.

### Phase locking and TWCO parameters

To test interactions between frontal theta and parietal alpha rhythms based on TWCO, we first estimated phase response curves (PRCs) for each participant and condition (see a representative participant in **Figure 5A**). Next, we quantified two key parameters of oscillator coupling: coupling strength and detuning (i.e., intrinsic frequency difference). Coupling strength was significantly reduced during the working-memory delay relative to fixation (*t*(25) = -2.86, *p* = 0.008, *d* = 0.56), and detuning was significantly increased (*t*(25) = 3.76, *p* < .001, *d*= 0.73) (**Figure 5C-D**). Together these changes make synchronization less likely during the working-memory delay. Accordingly, PLV was significantly lower during the working-memory delay compared to fixation (*t*(25) = -3.53, *p* = .001, *d* = 0.69) (**Figure 5B**).

**Figure 5.**
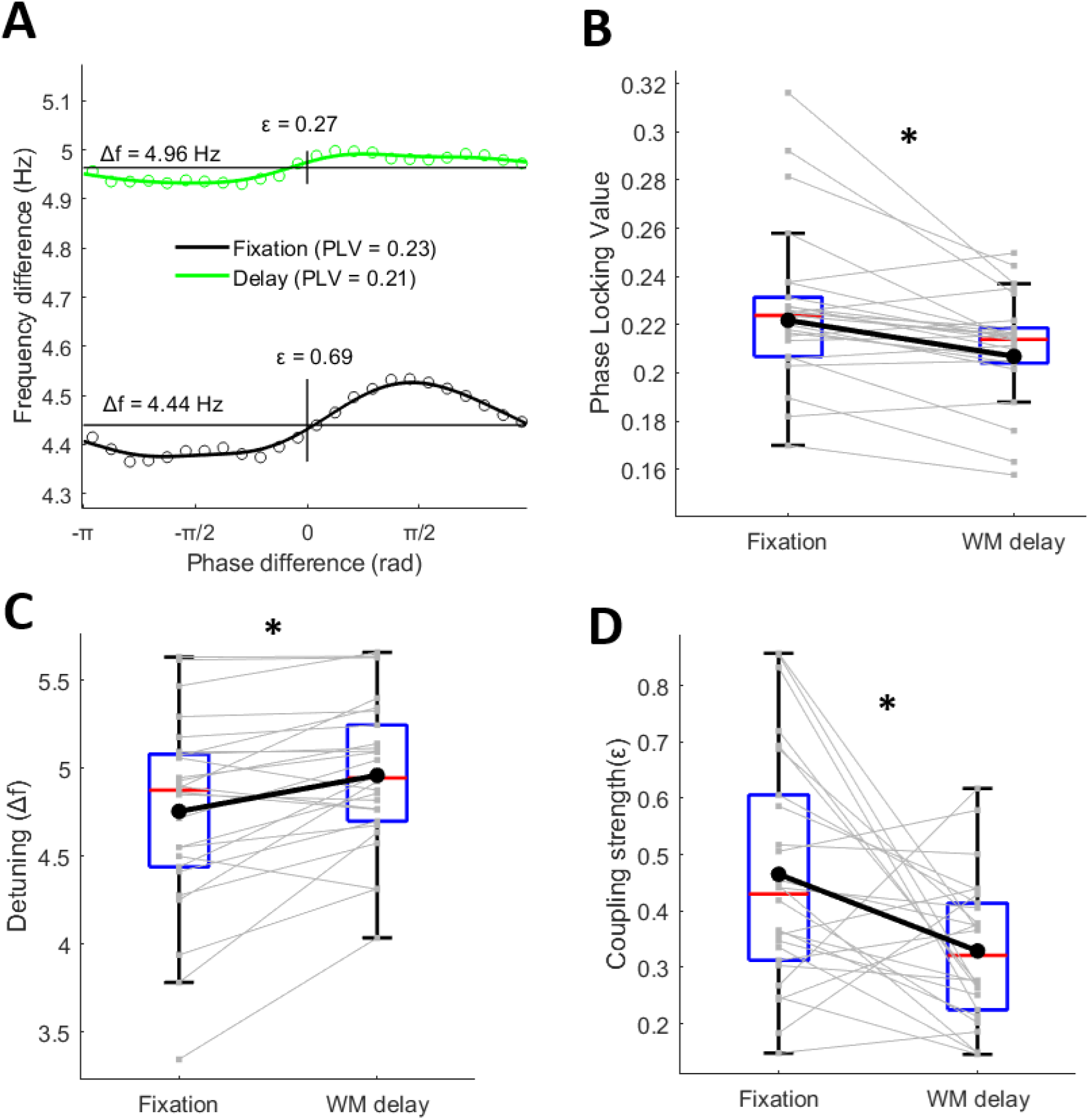
Modulations in phase locking and TWCO parameters. **A)** Phase Response Curve of a representative subject for fixation and working-memory delay. Instantaneous frequency differences between frontal theta and parietal alpha rhythms are plotted as a function of their phase differences. Detuning (Δf) was defined as the mean frequency difference across the PRC, while coupling strength was estimated as the peak-to-trough range of the fitted PRC (ε). **B)** Boxplots showing condition-related modulations in PLV, **C)** detuning and **D)** coupling strength. Individual lines reflect subjects and asterisks indicate significant differences at p<0.05.

### Relation to performance

To identify which neural metrics that best predicted behavioural performance, we conducted two stepwise linear regression analyses with accuracy and reaction time as the dependent variables and nine candidate predictors: PLV, detuning, coupling strength, alpha:theta mean ratio, parietal alpha frequency, frontal theta frequency, transfer entropy (anterior to posterior and posterior to anterior), and mutual information. While no significant predictors were identified for reaction time, PLV was a significant predictor of accuracy (*t*(25) = -2.40, *p* = .024). The regression coefficient for PLV was negative (β = –0.89, *SE* = .037; **Figure 6**), indicating that higher PLV values during the working-memory delay were associated with lower accuracy. None of the other predictors significantly contributed to the model.

**Figure 6.**
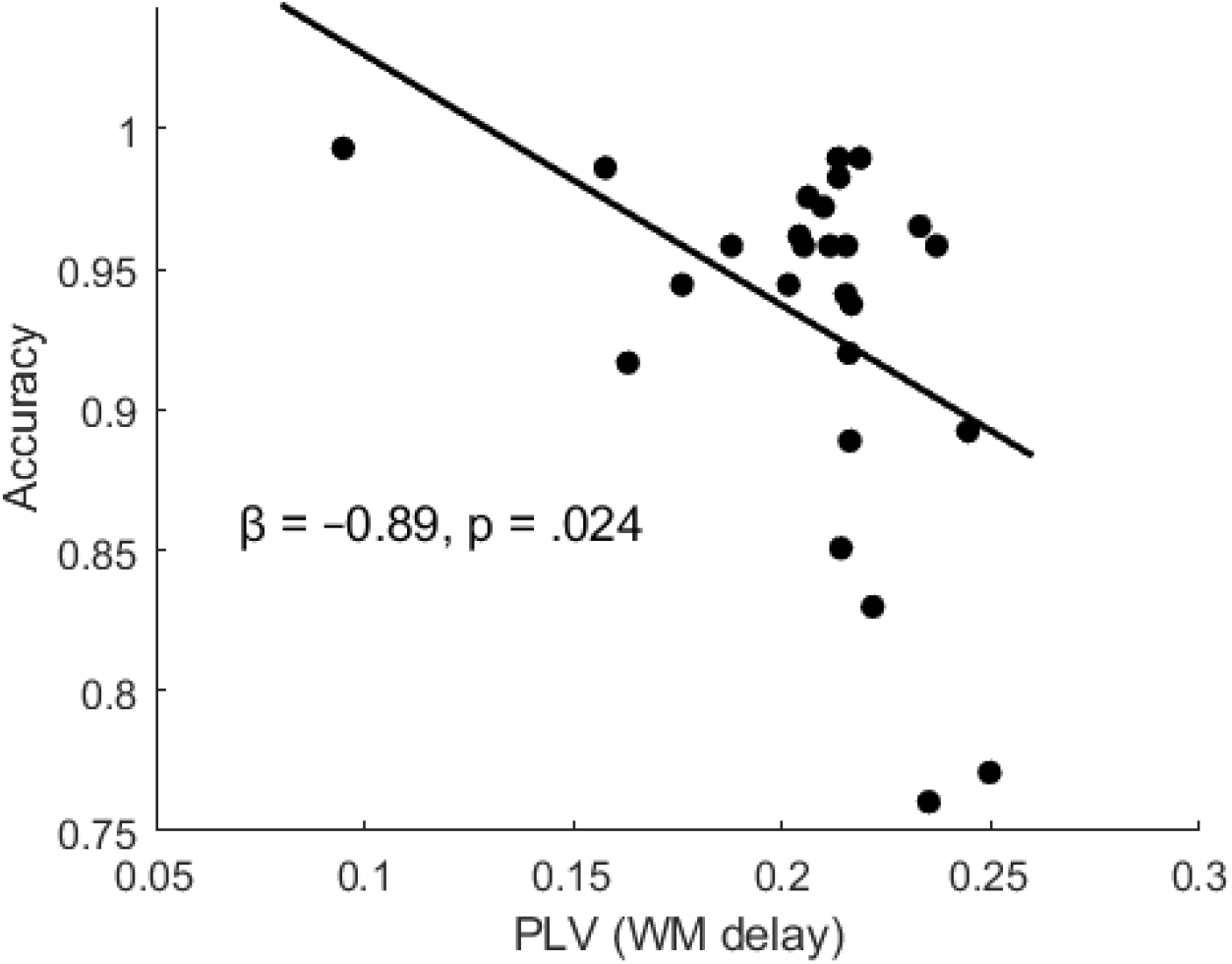
Relationship between PLV during the working-memory delay and behavioural accuracy. Each point represents one participant. A significant negative linear association was observed (*β* = – 0.89, *p* = .024), indicating that stronger phase locking between frontal theta and parietal alpha components was associated with lower task accuracy. Solid line shows the least-squares linear fit.

## Discussion

This study aimed to assess whether the previously reported cross-frequency dynamics between alpha and theta rhythms during working memory were accompanied by changes in inter-areal connectivity. For that purpose, we estimated both cross-frequency dynamics and connectivity, including model-free metrics from information theory, and model-based parameters derived from TWCO between brain sources generating EEG alpha and theta rhythms in the context of a working-memory task (N = 26). In line with previous literature, our results showed that working-memory retention (relative to pre-stimulus fixation) was associated with an increased occurrence of alpha:theta cross-frequency ratios around 2:1 and a decrease of ratios closer to 1:1. These cross-frequency modulations were due to a general increase in alpha frequency and decrease in theta frequency. The reported (cross)-frequency dynamics were accompanied by a decrease in inter-areal connectivity as quantified by Information theory metrics (i.e. reduced MI and TE) and TWCO parameters (reduced coupling strength and increased detuning). Finally, PLV (a standard measure of synchrony) was significantly reduced during working-memory delay and negatively correlated with behavioural performance. Together, our results suggest that the separation of frontal theta and parietal alpha rhythms in the frequency domain to form a ∼2:1 cross-frequency ratio during working-memory retention reflects functional segregation rather than functional connectivity between the respective neural generators.

According to the Binary Hierarchy Theory Brain Body Oscillations Theory (BHBBOT), the previously reported shift towards harmonic 2:1 alpha:theta cross-frequency configuration during working-memory retention (Rodriguez-Larios, Faber, et al., 2020; Rodriguez-Larios & Alaerts, 2019b) should reflect a relative connectivity increase between alpha and theta brain sources (Klimesch, 2018). This prediction was based on mathematical analysis showing that harmonic ratios between brain rhythms (e.g., 2:1, 3:1, 4:1) may lead to more stable excitatory phase meetings than non-harmonic ratios (such as the golden mean; i.e. 1.618…etc)(Pletzer et al., 2010). Contrary to this prediction we here show that the shift towards 2:1 alpha:theta ratios during working memory was accompanied by decreases (rather than increases) in connectivity between their brain sources, as quantified through information theory metrics. Our interpretation is that this is due to the relative decrease in cross-frequency ratios around 1:1 during working memory retention (**Figure 3**), which is the cross-frequency configuration that maximizes transient phase locking (Hoppensteadt & Izhikevich, 1998). Hence, our results suggest that areas oscillating at alpha and theta rhythms can increase their level of connectivity by transiently reducing their frequency ratio (instead of forming a harmonic cross-frequency ratio as suggested by the BHBBOT).

The Theory of Weakly Coupled Oscillators offers a theoretical framework to understand the here reported alpha:theta cross-frequency dynamics (Lowet et al., 2017, 2022). TWCO conceptualises synchronisation as a non-stationary and non-linear process in which modulations in frequency difference between pairs of oscillators allow coordination around a preferred phase-relation. Within this framework, phase synchrony is not maximised through fixed harmonic relationships (as in BHBBOT), but through transient reductions in frequency differences (i.e., detuning) that enable short-lived but functionally relevant periods of phase coupling. Our findings align with this view, showing rapid changes in frequency difference as a function of phase difference, characterized as the phase response curve. This suggests that during a cognitive task, the brain may favour flexible and dynamic coupling mechanisms based on frequency matching, as predicted by TWCO, rather than relying on fixed harmonic entrainment (as predicted by BHBBOT). By comparing the phase response curves between the fixation and working memory periods we could track what changes explained the reduction in connectivity during working memory retention. We found that the amplitude of the PRC reduced and the offset increased, indicating a reduction in coupling strength and an increase in frequency difference. This suggests that the brain makes use of both mechanisms to rapidly alter long-range functional connectivity between brain regions.

Given our research question, we limited the connectivity analysis to the main cortical generators of frontal theta and parietal alpha rhythms using data-driven spatial filters (Nikulin et al., 2011). In this regard, it is important to note that different networks (e.g., Default Mode Network, Salience Network, Dorsal Attention Network) have been shown to contribute to the generation of frontal theta and parietal alpha (Beldzik et al., 2022; Mantini et al., 2007; Marino et al., 2019; Ros et al., 2013; Zuure et al., 2020) and that this might have contributed to previous inconsistencies regarding EEG-derived fronto-parietal connectivity during working memory (Dai et al., 2017; Dimitriadis et al., 2016; Duma et al., 2019; Harding et al., 2015; Riddle et al., 2024; R. Wang et al., 2019). That is, it is possible that frontal theta and parietal alpha rhythms are generated by different networks (with different natural frequencies) during fixation vs. working-memory delay (Barzegaran et al., 2017; Rodriguez-Larios et al., 2022; Sokoliuk et al., 2019). In support of this idea, we show that the TWCO-derived parameter coupling strength, which has been previously linked to anatomical connectivity (Lowet et al., 2017), was significantly reduced from pre-stimulus fixation to working-memory delay, thereby suggesting the involvement of a different network. Nonetheless, it is also possible that coupling strength is modulated by other factors such as shared input or synaptic efficiency (Engel et al., 2021; Penn et al., 2016). Therefore, future studies combining EEG with other methods with greater spatial resolution (Fahimi Hnazaee et al., 2020; Iannaccone et al., 2015; Leicht et al., 2025) are needed to assess whether the generators of frontal theta and parietal alpha rhythms are different during fixation vs. working memory delay.

In summary, we show that during working-memory delay (relative to baseline) parietal alpha and frontal theta rhythms tend to separate in the frequency domain to form harmonic cross-frequency ratios around 2:1 (e.g. alpha = 12 Hz; theta = 6 Hz). Contrary to the predictions of a recent theory (BHBBOT), the increase in alpha:theta harmonicity during working memory delay was accompanied by decreases (rather than increases) in fronto-parietal connectivity as assessed though information theory metrics, TWCO parameters and phase synchrony. Moreover, phase synchrony between frontal theta and parietal alpha rhythms during working-memory delay was negatively associated with performance. Together, our results show that working-memory involves functional segregation between frontal regions generating theta rhythms and parietal regions generating alpha rhythms.

